# Low-cost 3D-printed inverted microscope to detect *Mycobacterium tuberculosis* in a MODS culture

**DOI:** 10.1101/2020.07.27.223701

**Authors:** Mario Salguedo, Guillermo Zarate, Robert H. Gilman, Germán Comina, Jorge Coronel, Patricia Sheen, Mirko Zimic

## Abstract

**Background:** The MODS is an important assay for early diagnosis of tuberculosis and drug susceptibility. MODS is based in the microscopic observation, underneath, of the characteristic cords of Mycobacterium tuberculosis colonies grown in liquid media. An inverted optical microscope is required to observe and interpret MODS cultures. Unfortunately, the cost of commercial inverted microscopes is not affordable in low resource settings in developing countries.

**Methodology:** To perform a diagnosis of tuberculosis using the MODS assay, images with modest quality are enough for proper interpretation. Therefore, the use of a high cost commercial inverted optical microscope is not indispensable. In this study, we designed a prototype of an optical inverted microscope created with a 3D printer and based on a smartphone. The system was evaluated by comparison of manual interpretations of 226 TB positive MODS culture images and 207 negative MODS culture images.

**Significance:** The prototype resulted in a low-cost inverted optical microscope, with simple functioning, and whose parts have been manufactured using 3D printing techniques. The quality of the images was good enough and achieved a 100% concordance between the manual inspection with the developed microscope, and the standard diagnostics of MODS.

## Introduction

Tuberculosis (TB) is still a global public health problem of high priority. Currently, an estimated 10 million people around the world have been infected with the disease and more than 1.2 million have died from it by 2019 [1]. Recently, cases of multidrug-resistant (MDR-) and extensively drug-resistant (XDR-) tuberculosis have been increasing, in part due to the lack of early diagnosis resulting in the absence of adequate treatment initiation [2]. Therefore, an assay for detection of MDR-TB and XDR-TB that is fast, simple and inexpensive is needed.

The Microscopic Observed Drug Susceptibility test (MODS) is the only single assay capable of detecting TB, determine MDR-TB and XDR-TB, directly from a sputum sample in just 7-10 days with high sensitivity and specificity [3, 4, 5].

MODS consists of the culture of a sputum sample in a liquid medium on a 24-well plate, which is incubated for 7-10 days. During the growth stage of bacteria present in the sample, they are suspended at the bottom of the well, allowing detection by microscopic observation of specific morphological cording patterns, which are characteristic of *M. tuberculosis*. This observation is done using an inverted microscope, which is a relatively expensive piece of equipment, which limits the use of this method in low resource settings.

In the last years, different prototypes of low-cost inverted microscopes have been developed as proofs of concept, which have been especially evaluated for the diagnosis of TB using the MODS assay [6, 7]. However, the complexity of some of these devices is still a disadvantage (use of heavy material such as aluminum and image acquisition system), in addition to a still considerable cost [6, 7].

In recent years, 3D printing (cast filament manufacturing (FFF)) has become a readily available consumer technology. Its ease of use and affordable cost, due to its mass production, has allowed its application in areas such as chemical instrumentation [8], microfluidics [9], biochemical analysis [10] and others. Recent studies have demonstrated the use of devices in the transmission of digital images for remote analysis by experts [11] or remote analysis using recognition software and artificial intelligence [12, 13].

Given the great growth of this technology and its various fields of application, in this study, we show the construction and evaluation of a 3D printed inverted microscope capable of capturing images of MODS cultures using a smartphone for further interpretation and diagnostics of TB.

## Methodology

### Optical system and the basic principle of operation

The optical system of our 3D printed microscope is based on two mirrors placed at an angle of 45 degrees to the horizontal with their reflective faces facing each other. These are simple, low-cost standard mirrors. On one of them, an objective lens is placed to capture the image that collects the light from the sample and on the other mirror, an ocular lens is placed, achieving a total magnification determined by the characteristics of both lenses. The distance between the lenses (optical distance) is approximately 120 mm, a little less than that recommended by the DIN standard (160 mm). The light enters vertically on the objective lens, is reflected from the first mirror horizontally to the second mirror and is again reflected vertically through the eyepiece lens. Finally, a smartphone is positioned in such a way that the camera’s sensor lens is aligned with the ocular lens. In this way, the smartphone serves to display and capture the images. Figure 1 shows an outline of this optical principle and its operation.

**Figure 1:**
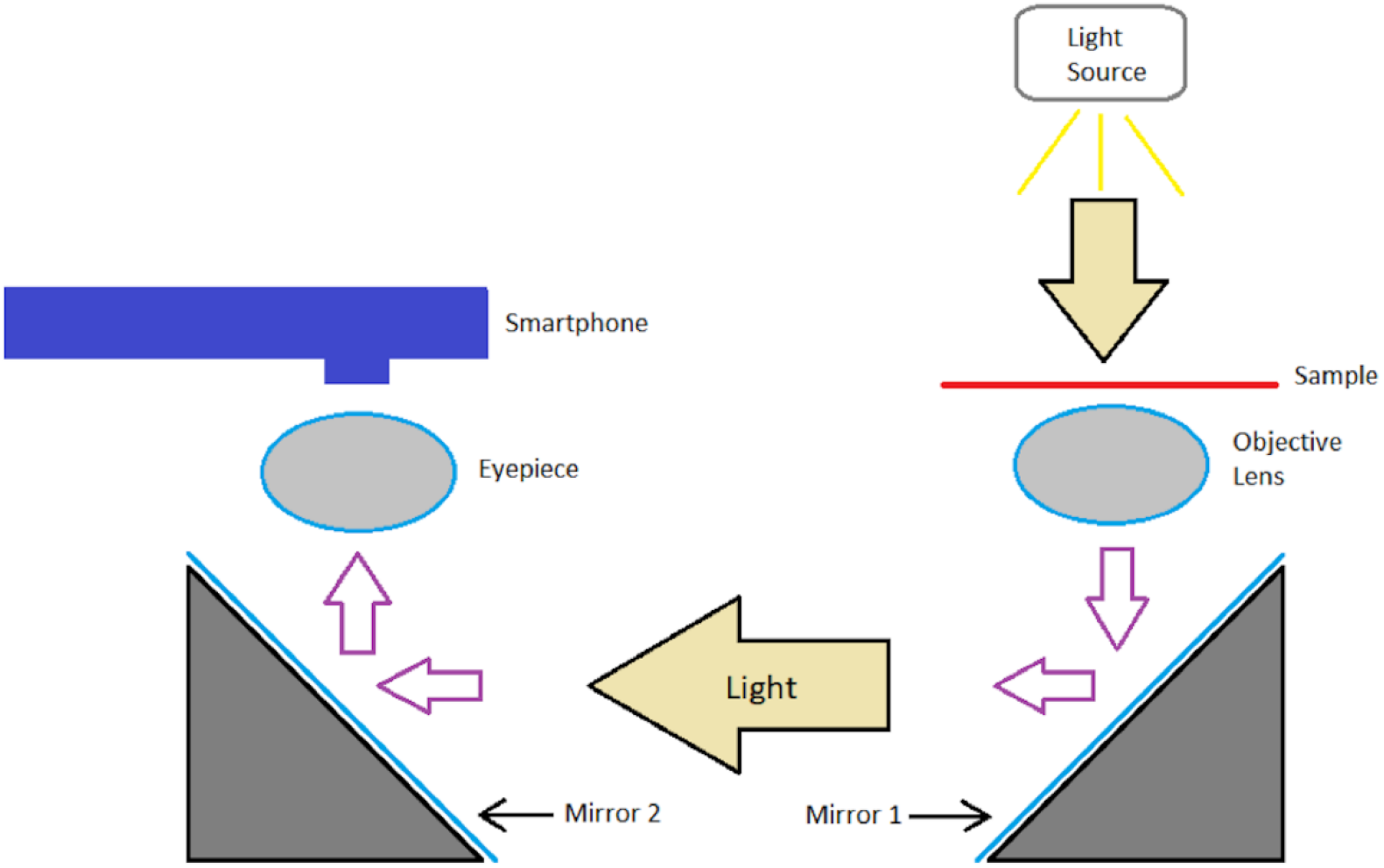
Working principle. The light from the source is transmitted through the sample, passes through the objective lens and is reflected on the mirror 1 horizontally and then on the mirror 2 vertically. It then passes through the eye lens and reaches the Smartphone’s camera sensor where the image is displayed.

### Prototype

The development of the prototype is based on the optical system (Figures 2a and 2b). This prototype consists of 19 pieces designed using CAD software. CAD files are saved in STL format, the same transformed into GCODE format using the Cura software. Each piece was 3D printed with an Ultimaker 2+ Extended printer. The printing parameters for the formation of the part in GCODE format (Cura Software) are printer extruder of 0.4 mm diameter, 0.2 mm thickness of each layer, 0.6 mm thickness in the first 3 and last 3 layers (total filling), 0.8 mm thickness in the lateral layers (total filling), 20% filling in the total part, printing speed: 60 mm/s, extruder travel speed: 120 mm/s. 2.85 mm thick Polylactic Acid (PLA) filaments from the same Ultimaker company were used.

**Figure 2:**
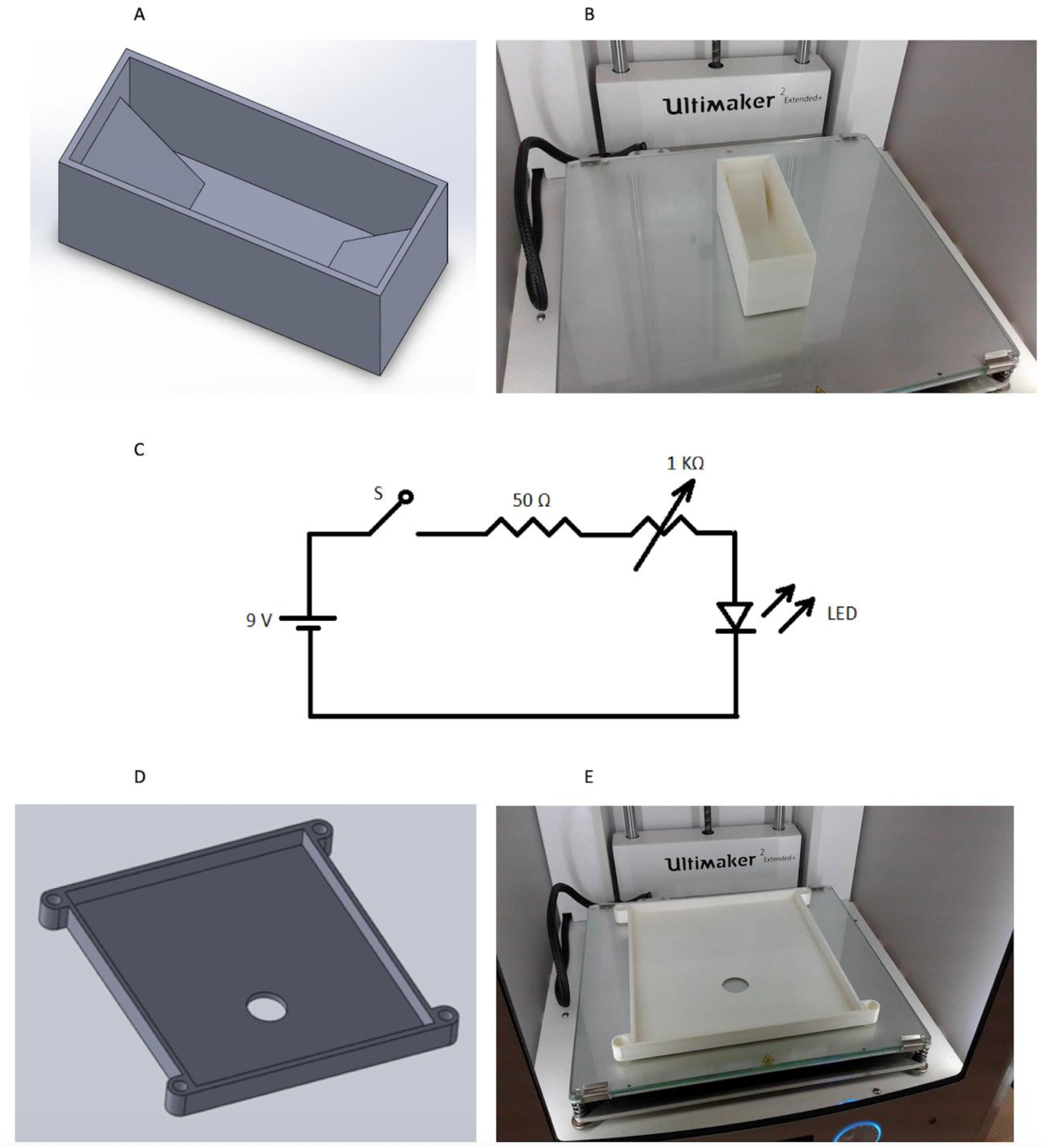
3D printed parts and lighting system circuit. (a) CAD software design of the mirror support. (b) Printed part of the holder. (c) Circuit diagram of the lighting system. (d) CAD software design of the platform for the plate support (MODS cultures). (e) Printed part of the plate holder.

The illumination system of the microscope is simple and adjustable, using a potentiometer and a white LED of 0.5 W power as a light source positioned 10 cm above the sample. The illumination circuit is powered by a 9V DC battery. The circuit design used is simple and easy to reproduce (Figure 2c). The microscope has a platform of a size to fit a 24-well MODS culture plate (Figure 2d and 2e). This platform has a manual positioning mechanism in the Z-axis that will serve to regulate the distance between the sample to be analyzed and the objective lens, and thus achieve the appropriate focus. The positioning mechanism consists of a 2mm pitch screw made in 3D printing and a fixed base that serves as a support for the screw (Figure 3a).

**Figure 3:**
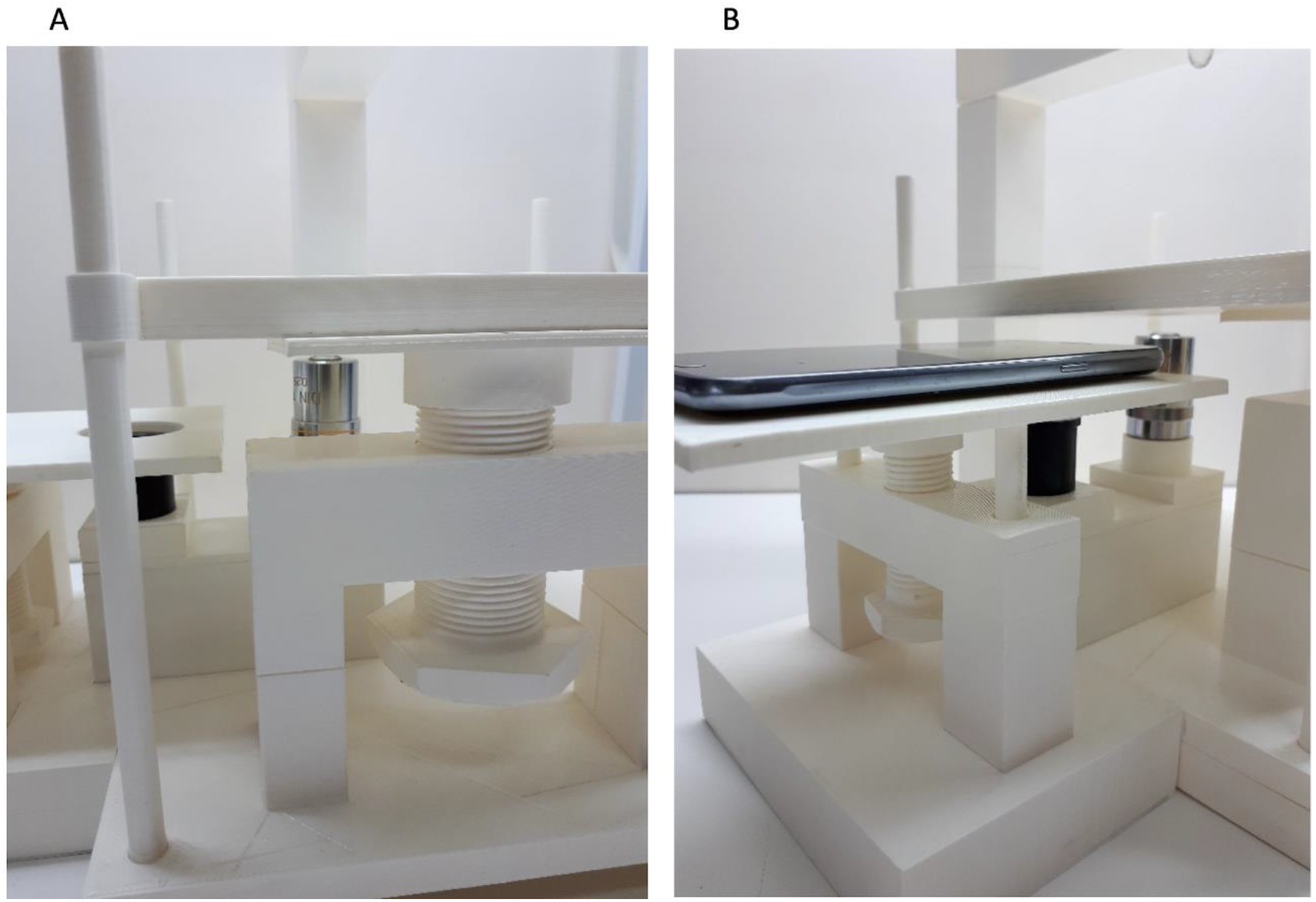
Z positioning mechanisms for focusing. (a) 2 mm pitch screw used for Z-positioning between the target lens and the plate. (b) 2 mm pitch screw used for Z positioning between the eyepiece lens and the Smartphone.

Above the other end of the optical system (over the ocular lens), there is a platform that supports the smartphone for image collection. This platform also has a Z-axis positioning mechanism for adjusting the distance between the smartphone lens and the eyepiece. Similarly, this Z-axis focusing system is given by a screw of 2mm pitch made in 3D printing and a fixed base that serves as a support (Figure 3b). Two standard low-cost microscope lenses were used for optical magnification. A 10X objective lens and a 10X eyepiece giving a combined magnification of 100X. A Samsung Grand Neo Plus smartphone, with main camera of 5 MP resolution, and Android 4.4.4 operating system (Kitkat), was used to view and capture the images.

The approximate cost to build this microscope is between $70 and $100 (not considering the cost of the smartphone), which is notably less than the cost of a commercial inverted microscope. The CAD design of the microscope (Figure 4a) and the files corresponding to the elaboration of each piece are available in the <u>web site</u>, as well as a <u>video tutorial</u> about the assembly upon request to the authors. The final microscope is shown as a lightweight structure of small size, which is quite portable (Figure 4b). The quality of the images taken from 9-day MODS cultures show the details necessary for correct interpretation (Figure 4c).

**Figure 4:**
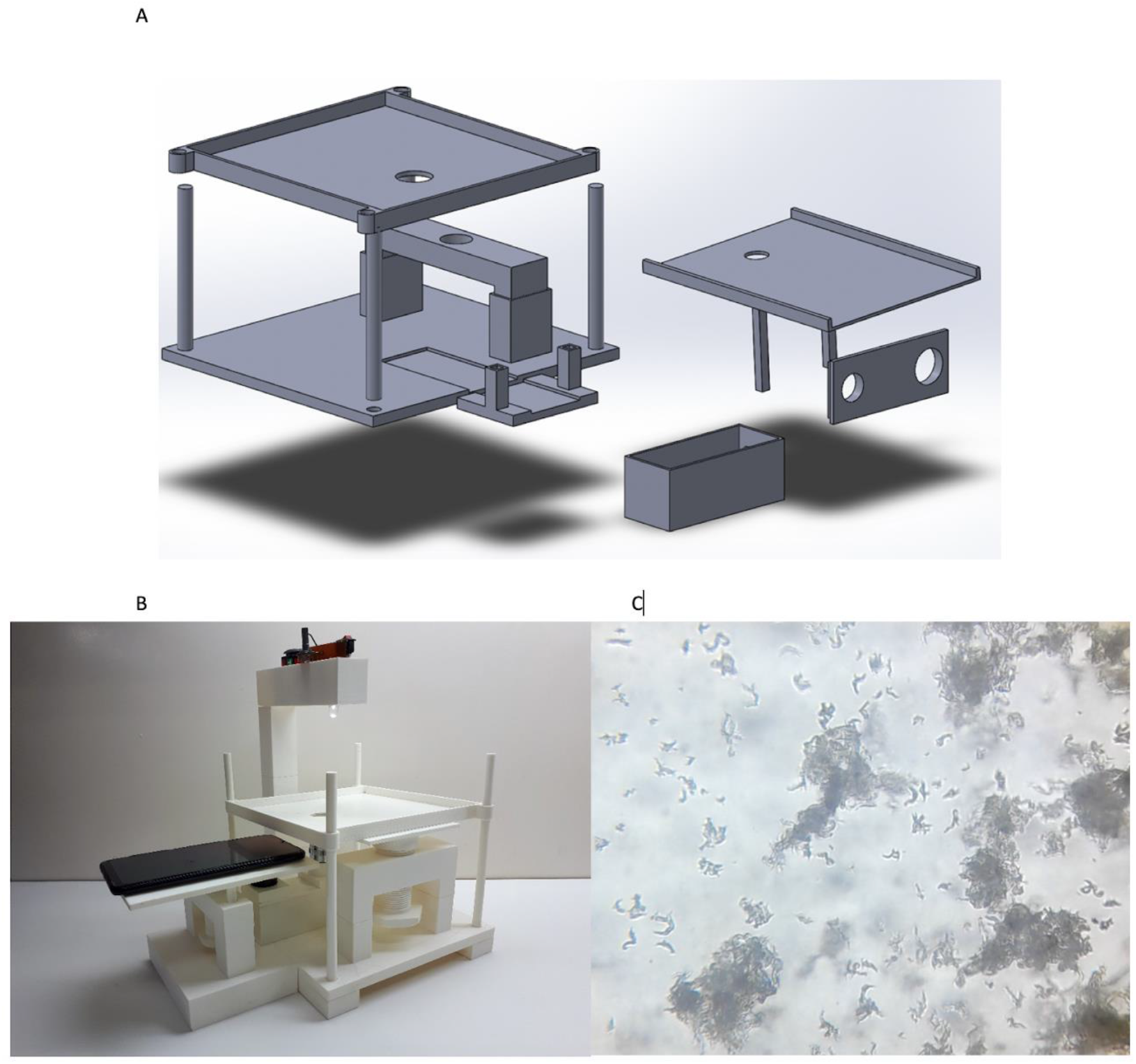
(a) CAD software design of some parts used in the inverted microscope assembly. (b) Inverted printed 3D microscope assembly (c) Image of a 9-day MODS culture taken with the inverted printed 3D microscope.

### Validation

The validation of the prototype was carried out by digitizing images from 20 different MODS cultures (10 positive cultures taken at 7 days and 9 days of incubation and 10 negative cultures). Sputum samples from TB patients and negative controls, for which MODS cultures were performed, were collected from surplus samples from another study. All images were captured with the Samsung Grand Neo Plus smartphone. A total of 114 images from 7 days positive cultures, 112 images from 9 days positive cultures and 207 images from negative cultures were obtained. All the digital images corresponding to the cultures performed are available <u>here</u> upon request to the authors.

## Results

### Performance of the prototype system for local TB diagnosis

All images captured with our 3D printed inverted microscope were analyzed blindly by a microbiologist with experience in MODS culture interpretation. Examples of these images corresponding to 7-day, 9-day positive cultures, and negative cultures are shown in Figures 5a, 5b, and 5c.

**Figure 5:**
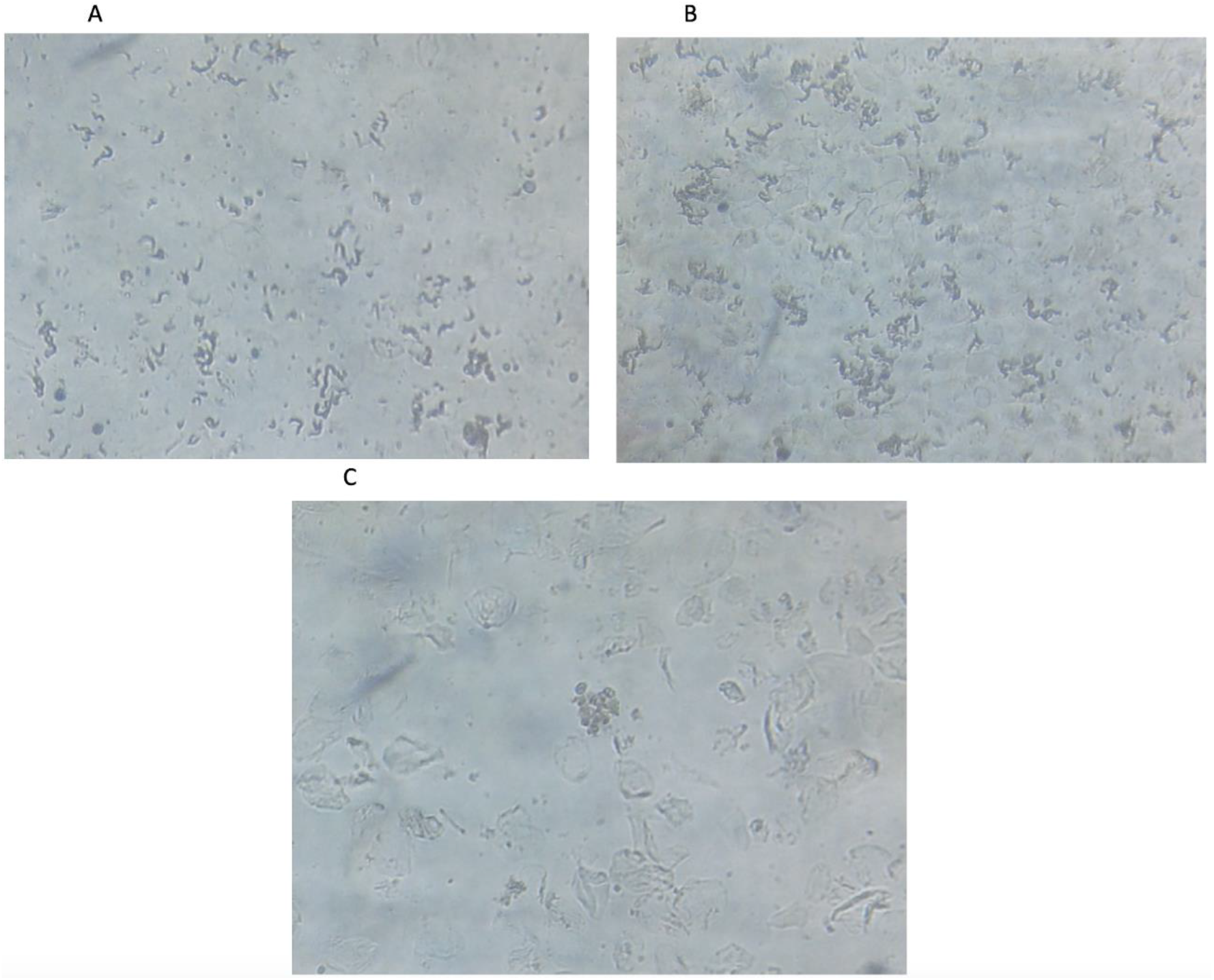
(a) Image of a 7-day MODS culture in the validation stage. The image shows the characteristic strands of Mycobacterium tuberculosis. (b) Image of 9-day MODS cultures in the validation stage. As well as the Mycobacterium tuberculosis cords, but with a bigger size and more defined clusters. (c) Image of a negative sputum sample.

Of the total of 226 positive images, 100% were confirmed by the expert as positive. Of the 207 images of negative cultures, 100% were confirmed as negative by the same expert. This evidence confirms that the quality of the images obtained through the 3D-printed built microscope is adequate to allow proper recognition and diagnosis of TB in MODS cultures.

## Discussion

The validity of the MODS assay depends on the quality of the microscopic images and the experience of the expert who reads and interprets them. Commercial inverted microscopes have usually been used to interpret MODS cultures. In many low-resource settings, it is virtually impossible to afford a commercial inverted microscope, resulting in a major obstacle to the application of MODS and limiting the early detection of TB cases as well as the early detection of MDR-TB and XDR-TB cases.

Our study demonstrates that high resolution and definition are not necessary for the correct interpretation of digital images of MODS cultures. It is demonstrated that through 3D printing technology we can build a very low-cost inverted microscope, accessible to the public, and capable of obtaining MODS culture images with sufficient quality for a correct analysis and interpretation.

The mass production of our methodology would produce even cheaper equipment and may be able to bring MODS to remote low resource settings. Previous studies have demonstrated the usefulness of smartphones in the transmission of digital images of MODS cultures for remote diagnosis [11], which is coupled with the development of a digital system and pattern recognition algorithms for automatic diagnosis using artificial intelligence tools with convolutional neural networks [13]. Both developments, together with the inverted microscope printed in 3D, provide a platform for the telediagnosis of tuberculosis from the analysis of digital images of MODS cultures in remote locations. In this way, our study contributes modestly to the global efforts to fight against tuberculosis.

